# Early detection of retinal alteration by visible and near-infrared optical coherence tomography (vnOCT) in a dexamethasone-induced ocular hypertension mouse model

**DOI:** 10.1101/478719

**Authors:** Weiye Song, Sipei Fu, Shangshang Song, Sui Zhang, Lei Zhang, Steven Ness, Manishi Desai, Ji Yi

## Abstract

**Purpose:** To apply a novel visible and near-infrared optical coherence tomography (vnOCT) in the dexa-methasone-induced ocular hypertension mouse model, and test the capability of four optical markers, peri-papillary retinal nerve fiber layer (RNFL) thickness, total retinal blood flow, VN ratio and hemoglobin oxygen saturation (sO_2_), in detecting retinal ganglion cell (RGC) damage in association with ocular hypertension.

**Methods:** Twelve mice (C57BL/6J) were separated into a control (n=6) and a dexamethasone group (n=6) receiving twice daily saline or dexamethasone eye drops, respectively, for 7 weeks. Intraocular pressure (IOP) measurements were taken at baseline and weekly. Optical measurements by vnOCT were longitudinally taken at baseline, 4 weeks and 7 weeks. Following week 7, *ex vivo* RGC counting was performed by immunostaining.

**Results:** The dexamethasone group showed a measurable rise in IOP by week 2. Despite the IOP differences between the dexamethasone and control groups, there was not a statistical difference in RNFL thickness or total blood flow over 7 weeks. The dexamethasone group did show an increase in retinal arteriovenous sO_2_ difference (A-V sO_2_) that was significant at week 4 and 7. The RNFL VN ratio showed a significant decrease at week 4 and 7 in dexamethasone group associated with a decreased RGC count.

**Conclusions:** RNFL VN ratio and A-V sO_2_ are capable of detecting early retinal alterations in the dexamethasone-induced ocular hypertension mouse model. Data analysis suggests VN ratio and A-V sO_2_ are corralated with RGC loss secondary to ocular hypertension, while being independent of IOP.

## 1. INTRODUCTION

Glaucoma is an irreversible progressive optic neuropathy associated with loss of retinal ganglion cells (RGCs)^1^. It is known as “the silent thief of sight”, since there are usually no symptoms until vision impairment has occurred and becomes irreversible. Typically by the time when functional changes are detected, 30% of RGCs have already been lost^2, 3^. As early treatment can significantly reduce the risk of glaucoma, preclinical detection prior to onset of functional visual field defects holds the key for prevention of vision loss^4, 5^.

Optical coherence tomography (OCT) has become the standard-of-care for glaucoma, and plays important role in diagnosis by identifying thinning in the peripapillary retinal nerve fiber layer (RNFL)^6-8^. The RNFL structural thinning is subsequent to RGC atrophy and denotes onset of glaucoma. However, RNFL thinning alone is not sensitive enough to differentiate early glaucoma from the high-risk glaucoma suspect who may benefit from earlier treatment^9^. Recent advance on optical coherence tomography angiography (OCTA) turns the attention to the microvasculature of the peripapillary region, by quantifying markers of the capillary flow index, and capillary density^10-15^. The sensitivity of those markers for early detection of glaucoma before the vision defect is still under investigation. While OCTA may prove to be valuable, the development of visible light OCT (vis-OCT) has recently drawn interest in the ophthalmic applications for several distinct advantages^16-18^. First, this technology allows functional measurement of hemoglobin oxygen saturation (sO_2_) as there is strong hemoglobin absorption in visible light range and by extension allows calculation of the inner retina oxygen metabolic rate (MRO_2_) when combining blood flow measurements^19^. Second, by comparing the reflected signal between visible light and near-infrared (NIR) OCT images, the light scattering spectroscopic (LSS) contrast can detect ultrastructural changes in the retina, on a length scale beyond the resolution limit of current commercial imaging devices ^20-22^. This LSS contrast can be quantified by a novel optical marker called the VN ratio that normalizes the visible light OCT images with NIR ones. Pathologic changes in the RGC cytoskeleton occur at a length scale <300 nm and use of LSS contrast or VN ratio can allow us to optically evaluate for changes prior to RNFL thinning^23, 24^. Lastly, vis-OCT can achieve 1-2 microns axial resolution^17, 25^, which could be beneficial to detect subtle RNFL thickness changes and refine current measurement capabilities.

The motivation of this study is to apply visible light OCT in a dexamethasone-induced ocular hypertension mouse model, as an early glaucoma model, and explore whether the spectroscopic markers can exhibit detectable changes. To fully utilize the advantage of vis-OCT, we have developed and implemented a novel fiber-based visible and near-infrared OCT (vnOCT) for simultaneous dual-band retinal imaging^21^. The combination of the two bands allows the multi-metric measurements of abovementioned spectroscopic markers, including VN ratio in RNFL to detect the ultrastructural changes, and sO_2_ in the inner retinal circulation. We characterized the model by IOP and RGC loss as compared to controls and longitudinally measured sO_2_, retinal blood flow, RNFL thickness, and VN ratio. The alterations and relation of these metrics may provide valuable insight into the pressure-related pathophysiology of glaucoma with use of vnOCT and lead to the development of novel optical markers that can translate into early detection of glaucoma clinically.

## 2. METHODS

All animal procedures were approved by the Institutional Animal Care and Use Committee at Boston Medical Center and conformed to the guidelines on the Use of Animals from the NIH and the ARVO Statement for the Use of Animals in Ophthalmic and Vision Research.

### Visible and near infrared optical coherence tomography (vnOCT) setup

The dual band OCT setup used in this study is modified from our previous publication^21^, as shown in Fig. 1. Briefly, a supercontinuum laser provided both visible light and NIR illumination, from 540-600 nm and 800-900 nm respectively. The visible light was separated by a dichroic mirror (DMLP650R, Thorlabs), dispersed by a pair of prisms, and filtered by a slit aperture. The filtered light was reflected by a mirror and coupled in an optical fiber. The NIR range was also separated by another dichroic mirror (DMLP900R, Thorlabs), filtered by two edge filters (DMLP805, Thorlabs and 19354, Edmund optics), and coupled into an optical fiber. Two bands were combined by custom-made wavelength division multiplexer (WDM), and directed into a 90/10 optical fiber coupler (TW670R2A2, Thorlabs). The returning light was separated by another WDM for visible and NIR channel, and recorded by two separate spectrometers. Each spectrometer is equipped with a 2048 pixel line scan camera (Basler, splk2048-140k). The spectrometers were running at 50 kHz A-line rate. Because both bands propagate within the same optical fiber, they shared the same sample arm and reference arm. The sample arm consisted of a collimator (*f* = 6mm), a two axis galvanometer scanner (GVSM002, Thorlabs), and a 3:1 telescope (*f*=75mm and *f*=25mm) to relay light to the pupil. The reference arm consisted of a collimator (*f*=10mm), several dispersion control BK-7 plates, a variable neutral density filter, and a mirror to reflect the light. The beam size on the pupil is about 0.2mm in diameter. The incident optical power on the mouse pupil is 0.3 mW and 0.8 mW, in visible light and NIR bands, respectively. The system acquires visible and NIR OCT images simultaneously.

### *In vivo* retinal imaging

For imaging, we anesthetized mice with a mixture of ketamine and xylazine (ketamine: 11.45 mg mL^−1^; xylazine: 1.7 mg mL^−1^, in saline) that was injected intraperitoneally. The dose was 10 mL per kg body weight. After anesthesia, the animal was placed on a custom-made animal holder for imaging. We applied 1% Tropicamide Hydrochloride ophthalmic solution to dilate the pupil. Commercial artificial tears were applied to the mice’s eyes every other minute to prevent corneal dehydration. The imaging included two scanning protocols consecutively performed, including three-dimensional (3D) OCT images by a 256 × 256 raster scanning pattern centered at the optic disk, followed by a dual-circle scanning that scanned two concentric circles with 4096 pixels in each circle with each circle repeated 8 times. The optic nerve head (ONH) was aligned to be at the center of the field-of-view (FOV) so that all the major vessels could be imaged. Figure 1A-1B exemplify the *en face* projection of visible and NIR OCT images, acquired at the same time. The 3D OCT images were used for calculating sO_2_, and VN ratio. The dual circle scanning data was used to calculate retinal blood flow and RNFL thickness. After imaging, the mice were released from the animal holder for recovery.

**Fig. 1.**
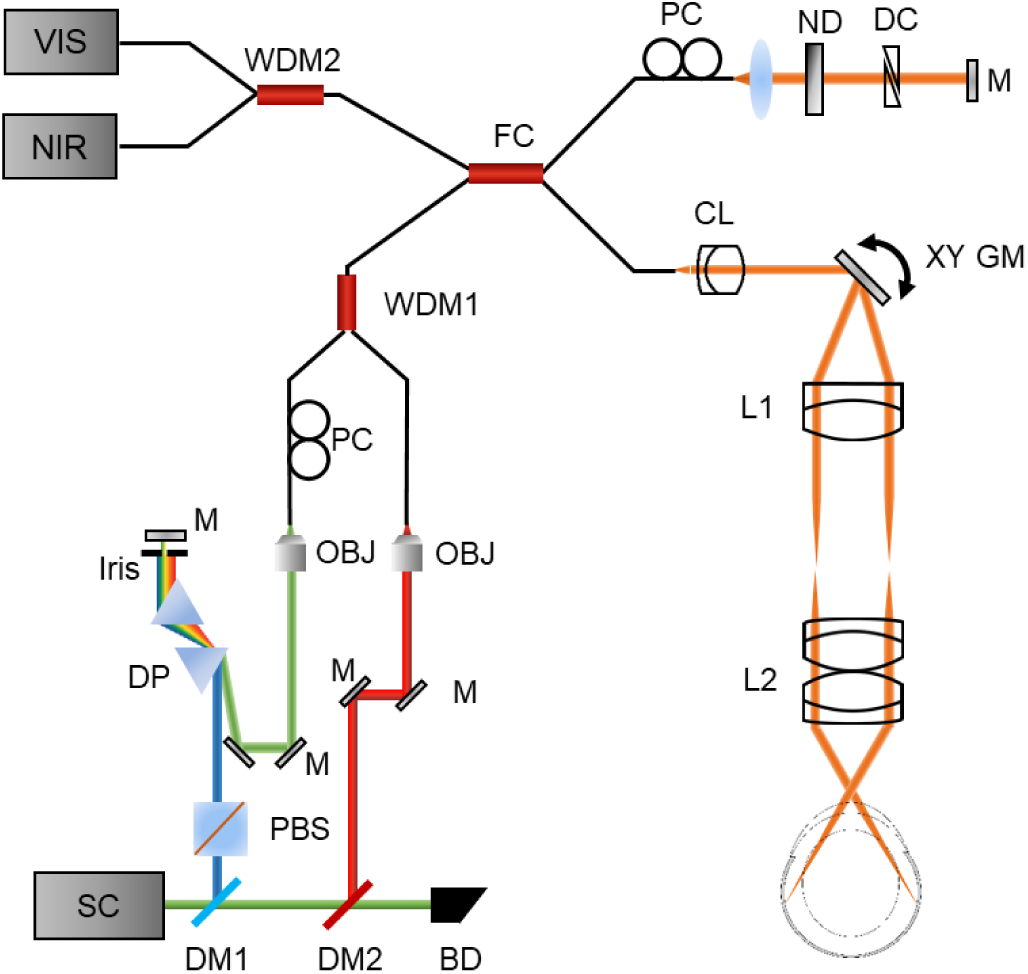
Schematic of the vnOCT system setup. Two wavelength division multiplexers (WDM) and a fiber coupler (FC) merges and split the visible and NIR channel. The same sample and reference arms are shared by two channels. These two channels were collected by separate spectrometers. SC: supercontinuum source; DM: dichroic mirror; BD: beam dumper; PBS: polarization beam splitter; DP: dispersive prism; M: Mirror; OBJ: objective lens; PC: polarization controller; CL: collimating lens; GM: galvanometer mirror; L: lens; ND: neutral density filter; DC: dispersion compensation plates.

### Blood hemoglobin saturation (sO_2_)

We used the previously published method for calculating sO2 from retinal arteries and veins ^16,19^, as illustrated in Fig. 2C. The 3D images from the visible light channel were used. A short time Fourier transform with a sweeping Gaussian window (FWHM *kw = 0.32 μm*^*-1*^) was performed to generate wavelength-dependent vis-OCT images. We used 11 windows to generate four-dimensional data (*x, y, z, λ*). The retinal surface was identified by setting an intensity threshold. To reduce the dimension, individual vessels were first segmented from the *en face* projection, and all the A-lines within each vessel were lined-up to a 2D B-scan. Then each A-line was shifted in reference to the retinal surface. The signal from the vessel bottom wall of each individual vessels was averaged, and the same process was iterated for each spectral window to generate a spectrum for each vessel. Lastly the following analytical model was used to fit the spectrum to calculate sO_2_ from 545-580 nm:

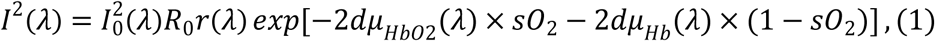

where *I*_*0*_ *(λ)* is the incident spectrum on the retina; *R*_*0*_ is the reference arm reflectance; *d* [mm] is the vessel diameter; *r(λ)* [dimensionless] is the reflectance at the vessel wall, whose scattering spectrum is modeled as a power law under the first-order Born approximation *r(−λ)* = *Aλ*^α^, where A [dimensionless] is a constant26. The subscripts Hb and HbO2 denote the contribution from deoxygenated and oxygenated blood, respectively. The optical attenuation coefficient μ [mm^−1^] combines the absorption (μ_a_) and scattering coefficients (μ_s_) of the whole blood, which are both wavelength- and oxygenation-dependent

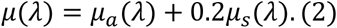

**Fig. 2.**
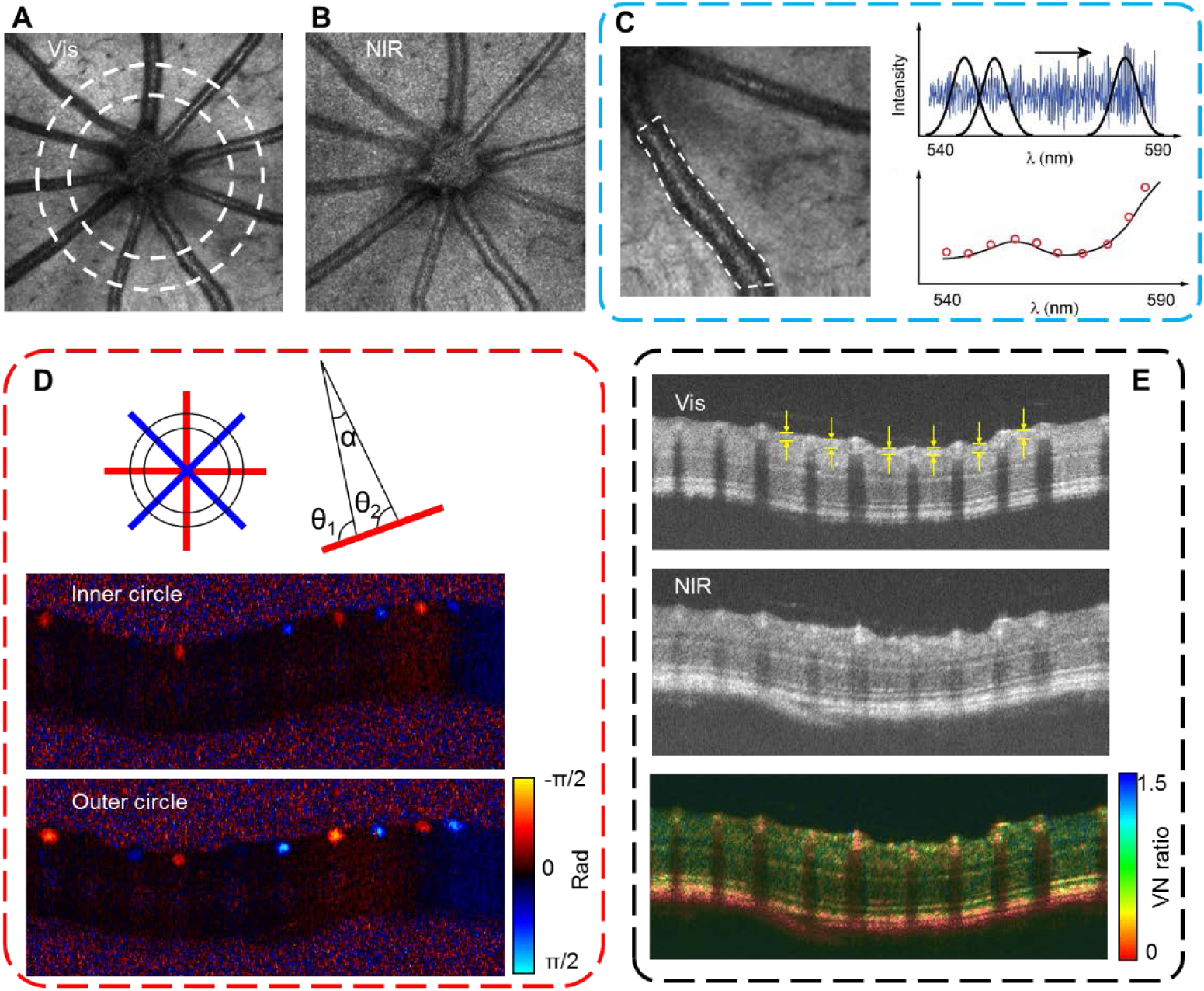
Multi-metric measurements by vnOCT. (A-B) En face projection of mouse retina in visible and NIR channel, respectively. Two concentric circles represent the dual circle scanning pattern. (C) Method for sO2 calculation by segmenting the vessel and the subsequent spectroscopic analysis. Vessel location was first manually segmented, and Short Time Fourier transform was performed to obtain 4D data (x,y,z,λ). Finally, the spectra from bottom vessel wall was averaged and fitted by Equation 1. (D) Illustration of RNFL thickness and VN ratio calculation. The circular B-scan image in visible OCT was used to calculate RNFL thickness, as shown in yellow arrows. The intensity ratio between visible and NIR channels was defined as VN ratio. (E) The schematic of blood flow measurements. The difference of the phase contrast from inner and outer circle was used to calculate Doppler angle and blood velocity.

The value of µ_a_ and µ_s_ is provided by literature ^27^. A least-square linear regression to the logarithmic spectra returned the value of sO_2_. We averaged sO_2_ from all the arteries and veins, respectively for each retina. One example of the sO_2_ calculation is shown in Supplemental Fig. 1.

### Retinal blood flow

We used the dual-circle scanning data from NIR channel to calculate the blood flow, using the method of phase sensitive Doppler OCT (Fig. 2D) ^28-30^. The scanning protocol scanned two concentric circles around the ONH at a radius of 0.21, and 0.32 mm, respectively ^31-33^. At each circle, it first scanned clockwise and immediately counter clockwise, and the round trip was repeated 4 times. Each circular scan included 4096 A-lines. The phase delay between adjacent A-lines can measures the blood flow velocity, *v* (mm/s) by

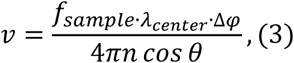

where *f*_*sample*_ ^34^ is the OCT A-line scanning frequency; λ_*center*_ [nm] is the center wavelength of the OCT light source at 840 nm; δ φ [rad] is the phase shift between adjacent A-scans; n = 1.38 [dimensionless] is the refraction index of the sample. The Doppler angle *θ* [rad] is the angle between the probing beam and the vessel.

The only unknown parameter in the right side of equation is the Doppler angle θ. To estimate *θ*, we used the phase difference between two circles^35^, as illustrated in Fig. 2D. The blood flow velocity from the same vessel measured at the inner circle *v*_1_, and the outer circle *v*_2_, should be the same such that *v*_1_ *= v*_2_. When plugging Equation 3, we have

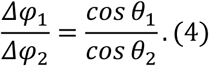

The difference between *θ*_1_ and *θ*_2_ is controlled by the galvanometer scanner that *θ*_1_ = *θ*_2_ +α, where α = 0.035 [rad]. Therefore, by taking the ratio of the phase difference between two circles from the same vessel,

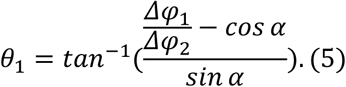

The data processing follows several steps. First, the phase delays between adjacent A-lines were calculated after the Fourier transform. Second, the phase signals from the clockwise and counter clock wise circular scanning were averaged, after reverting the counter clock wise scan. The purpose of the round trip scanning is to cancel out the phase background due to the retinal surface topography. Then the phase signals within each vessel were averaged for *δφ*, after manually segmenting the vessel lumens. Four repeated measurements at each circle were averaged. The Doppler angles were calculated according to Equation 5, and so were the velocities according to Equation 3. Finally, the blood flow F [µL/min] was calculated by:

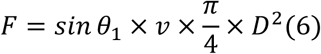

where *D* [µm] is the diameter of the vessel which was calculated by the vessel lumen area segmented manually.

### RNFL thickness measurements

Inner circular scanning data from visible channel was used for measuring RNFL thickness, for better axial resolution. We used circular scanning as it has much higher sampling density, better signal-to-noise ratio and is also consistent with the clinical protocol. Fig. 2E shows the identification of RNFL in B-scan images. We used Image J to manually measure the RNFL thickness. We used multiple measurements (≥5) at different radial locations and then averaged them to obtain the value for each retina.

### Spectroscopic marker VN ratio

The method to calculate VN ratio has been described in our previous publication^21^. The VN ratio quantifies the spectroscopic contrast between the visible and NIR channels, and thus allows ultrastructural measurements beyond the resolution limit of OCT. The VN ratio is calculated based on the absolute image intensity of OCT images in the visible and NIR channels. The noise floor calculated above the retina in the image was subtracted in both the visible and NIR channels.

To calculate VN ratio, we first interpolated the NIR cross sectional (B-scan) image in the axial dimension for a consistent scale to the visible B-scan image, and co-registered the two images by shifting the NIR B-scan image. The surface of the inner retina was detected by setting an intensity threshold. The intensity from the superficial 30μm was integrated into both channels, primarily consisting of RNFL and RGCs; and the ratio between visible over NIR channel produced the VN ratio. We performed manual image segmentations on the *en face* OCT projections to select region of interest (ROI) and to avoid blood vessels (See supplemental Fig. 2 for an example of ROI selection). The VN ratios from all ROIs were averaged to represent the reading for one retina.

To ensure the measurements are free of systemic variation, we used the VN ratio from blood as an on-site reference *in vivo*. The VN ratio by integrating 50μm thick of whole blood is theoretically calculated to be ∼0.6 with <10% variation between oxygenated and deoxygenated blood^21^. Then, the VN ratio from RNFL/RGCs is adjusted by:

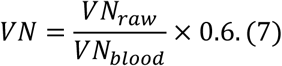

### Dexamethasone-induced ocular hypertension mouse model

An established protocol of dexamethasone-induced ocular hypertension mouse model^36-38^ was used in this study. Ten-weeks-old C57BL/6 male mice were purchased from Jackson lab. Intraocular pressure (IOP) elevation was achieved by treating the eyes with 0.1% dexamethasone eye drop twice daily for 7 weeks. The control groups received saline eye drops the same time as the IOP-elevated group. The IOP from each eye were measured weekly by a tonometer until the termination of this study. For eye drop administration and IOP measurements, mice were under isoflurane anesthesia. Specifically, animals were placed in the induction chamber for ∼2 mins under 3% isoflurane carried by 1.5L/min oxygen flow. Then the animal was placed on a stage, under 1.5% isoflurane was delivered by a nose cone. The IOP measurements were measured within 3 mins from the induction to avoid the influence of anesthesia on IOP.

### Eye dissection and Immunohistochemistry

At the end point of the study (7 weeks), mice were euthanized and their eyes were enucleated using a curved tip tweezer. Retinas were subsequently dissected, and fixed in -20°C methanol. For immunofluorescence staining, the retinas were first rinsed in PBS to wash away the methanol, and then were washed three times (10 minutes each) in 0.05% PBST (0.05% Triton X-100 PBS solution). After that, retinas were blocked/permeabilized for 1.5 hours at room temperature in a solution containing 0.25% Triton X-100 and 10% goat serum in PBS. Immunostaining was performed using primary and secondary antibodies. Primary antibody, rabbit anti-Brn3a (Millipore Sigma) was diluted 1:500 in 5% goat serum in PBS and incubated with retina overnight at 4°C. After 3 washes in 0.05% PBST, retina was incubated for additional 2 hours at room temperature with Alexa Fluor 488 conjugated goat anti-rabbit secondary antibody (Invitrogen) diluted (1:800) in 10% goat serum in PBS. Finally, the retinas were rinsed 2 times with PBS, mounted in aqueous mounting medium, and cover-slipped for imaging.

### Cell counting

After immunofluorescence staining, the whole mount retina slides were imaged by a confocal fluorescence microscope (Leica SP5, 10x Objective). The field-of-view of each image is 0.78^2^ mm^2^. The number of stained RGCs was manually extracted from the confocal images in Image J, within a ROI to avoid large vessels and non-uniformed locations subsequent to the process of immunostaining. The RGC cell density was then calculated by the cell number per ROI area. At least 4 ROIs from each retina image were used, and the average cell density representing the readings from each retina.

### Statistical Analysis

Two samples student’s T-test was used in Fig. 3 and 4. The Pearson’s linear regression was used to test the correlation among IOP, VN ratio, and sO_2_ in Fig. 5.

**Fig. 4.**
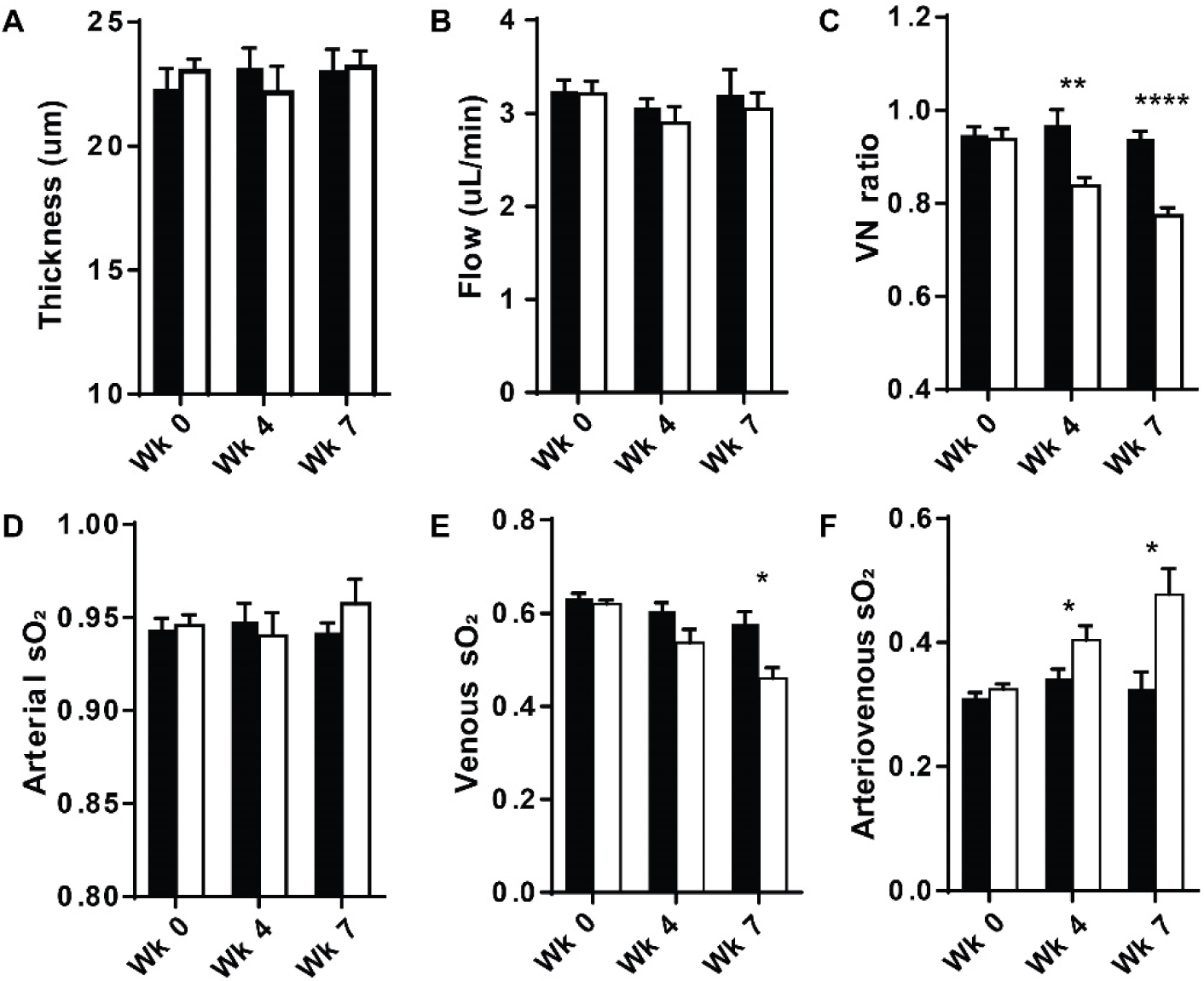
The longitudinal measurements by vnOCT at week 0, 4 and 7 for (A) NFL thickness (μm), (B) blood flow (μL/min), (C) VN ratio, (D) arterial sO2, (E) venous sO_2_, and (F) arteriovenous(A-V) sO2 difference. *P<0.05; **P<0.01; ****P<0.0001.

**Fig. 5.**
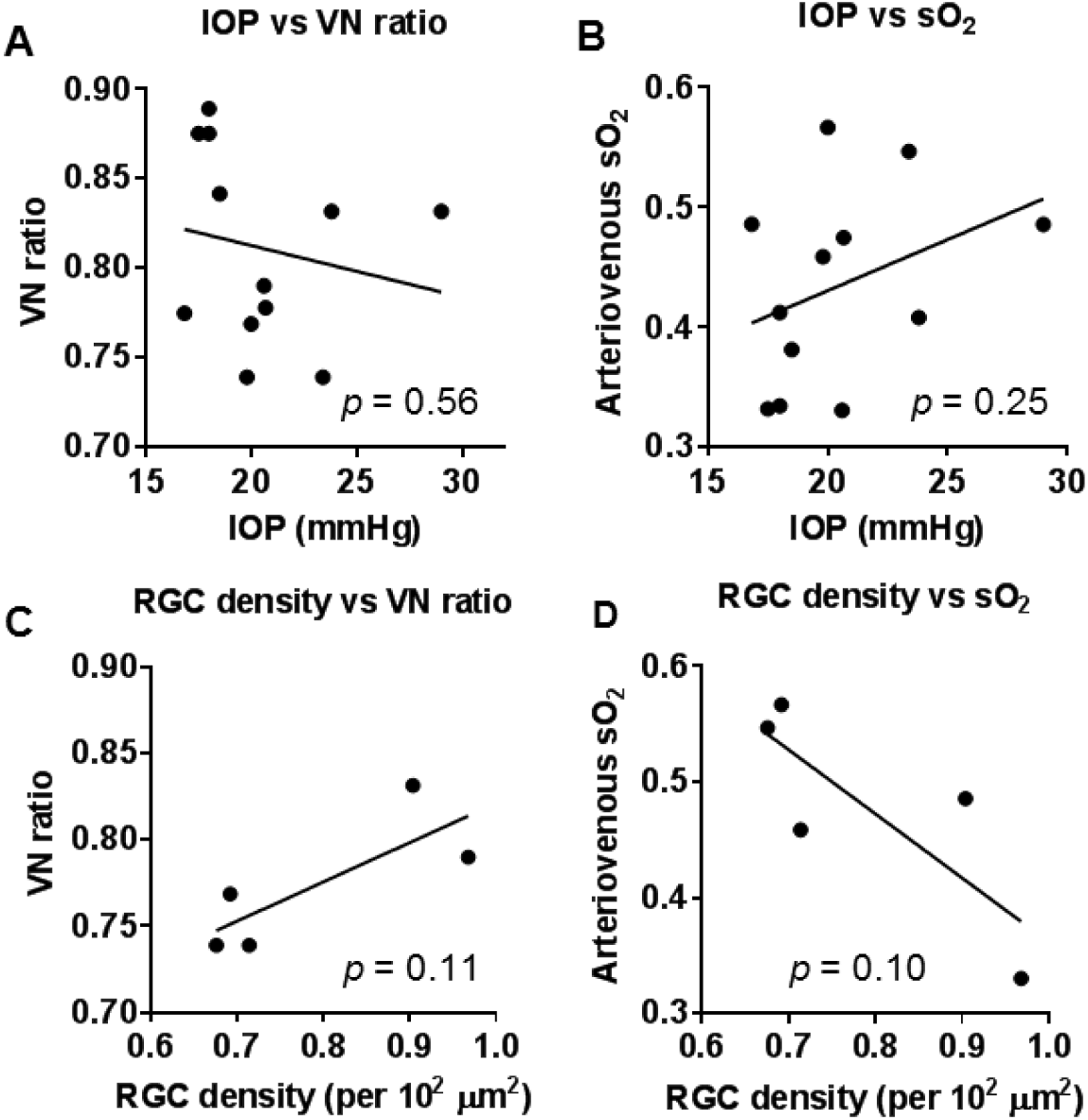
Correlation among vnOCT markers, IOP and RGC density. (A-B) The correlation between VN ratio, A-V sO2 difference and IOP on each eye in dexamethasone group at week 4, and 7. (C-D) The correlation between VN ratio, A-V sO2 difference and RGC density on each eye in dexamethasone group at terminal week 7.

**Fig. 3.**
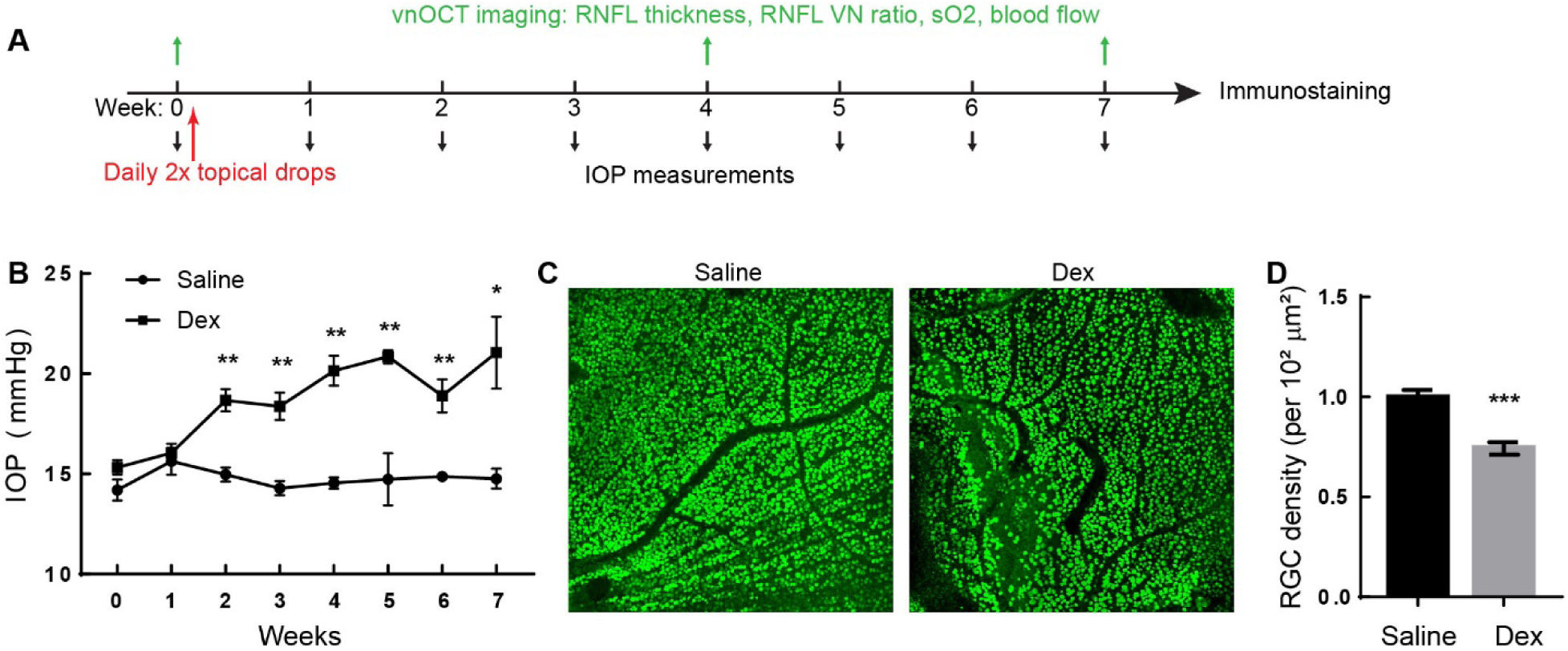
Characterization of the dexamethasone-induced ocular hypertension mouse model. (A) The timeline of the measurements over the span of 7 weeks. (B) Weekly measurements of intraocular pressure by a tonometer. (C-D) Representative confocal fluorescence microscopic images of anti-Brn3a immunostaining for RGCs, in saline and dexamethasone group. (E) The comparison of the RGC density between saline and dexamethasone groups. *p<0.05, **p<0.01, ***p<0.001.

## 3. RESULTS

The study design is illustrated in Fig. 3A. IOP measurements were performed weekly to confirm the induced ocular hypertension. The vnOCT imaging was performed longitudinally at the baseline week 0, and at week 4 and week 7. The study was terminated at 7 weeks of administration, and the effectiveness of the model was further verified by the RGCs loss via immunostaining and cell count. The study started with 12 animals, 6 in dexamethasone group and 6 in control group. Four animals from each groups reached the terminal stage at week 7.

The animal model was first validated by measuring IOP and RGCs loss. The baseline IOP was noted to be in the mid-teens for both control and dexamethasone group, respectively, and this was not statistically different. The IOP in the dexamethasone group increased from week 2 onward and this increase was statistically significant as compared to the control group (p <0.05, Fig 3B). The characteristics of the IOP change are consistent with multiple previous publications using this animal model^36-38^. The RGC density (cells per 10^2^ μm^2^) as measured by confocal fluorescence microscopic images (Fig. 3C, 3D) *ex vivo* was decreased in the dexamethasone group compared to the control group and this was statistically significant (p=0.0002, Fig. 3E), confirming the RGC loss.

The results of longitudinal multi-metrics measurements by vnOCT are summarized in Fig. 4. The RNFL thickness were similar at baseline between the two groups, and did not change significantly from baseline to week 4 or 7 for either group, or comparatively (Fig. 4A). Similarly, blood flow measurements showed no significant change between the dexamethasone and the control group (Fig. 4B). The blood flow did appear to have a moderate decrease comparing the dexamethasone to the control group at week 4 and 7. When we examined the VN ratio measurements, the values were similar at baseline between the two groups, but decreased progressively at week 4 (p=0.0048) and 7 (p=0.00009) in the dexamethasone group compared to the control group (Fig. 4C). This change is consistent with the previous reported spectroscopic change due to the damage of ultrastructural cytoskeleton loss in RGCs^24^.

The retinal arterial and venous sO_2_ were similar at baseline between the two groups as was the arteriovenous sO_2_ difference (A-V sO_2_) between the two groups. The arterial sO_2_ did not show any statistically significant change temporally or comparatively between groups throughout the study. The venous sO_2_ did show a progressive drop in the dexamethasone group compared to the control, which was statistically significant at week 7 (p=0.01, Fig. 4E). As a result, the A-V sO_2_ also showed a progressive rise at week 4 (p=0.039) and week 7 (p=0.016), indicating a higher oxygen extraction efficiency from the inner retinal circulation (Fig. 4F).

The vnOCT measurements showed that VN ratio and A-V sO_2_ were significantly altered in dexamethasone group. We next examine the correlation of these two markers to IOP and RGC loss within the dexamethasone group. We first compared VN ratio, and A-V sO_2_ with IOP at weeks 4 and 7 in ocular hypertensive eyes (Fig. 5A, 5B), and only weak correlation was found by Pearson’s correlation test. We next compared RGC density measurement with the terminal measurement of VN ratio and A-V sO_2_ difference at week 7 (Fig. 5C, 5D). Although the sample number is small, there appears to be moderate correlation between VN ratio, A-V sO_2_ and RGC loss. This analysis suggests that VN ratio and A-V sO_2_ may effectively serve as markers for RGC damage, while being independent of IOP.

## 4. DISCUSSION

In this study, we demonstrated that spectroscopic markers (VN ratio, A-V sO2) enabled by vnOCT have the potential to detect ultrastructural changes in the RNFL, as well as alterations in blood hemoglobin sO_2_ within the inner retinal circulation associated with ocular hypertension in a dexamethasone-induced ocular hypertension mouse model. These findings suggest that a decreased VN ratio and increased A-V sO2 difference may represent early changes in development of glaucoma.

The dexamethasone mouse is a well-established model of steroid induced ocular hypertension and glaucoma^36-38^. Consistent with previous reports, our mice developed significantly increased IOP within 2 weeks of starting topical dexamethasone drops with subsequent progression to RGC loss. The mechanism by which steroids cause ocular hypertension remains to be fully elucidated but decreased outflow facility is the main contributing factor. Phulke *et al.* has nicely outlined the leading molecular pathways to reduced outflow facility including accumulation of glycosaminoglycans, laminin, elastin and fibronectin in the trabecular meshwork (TM), increased TM cell nuclear size and DNA content, and decreased prostaglandin synthesis^39-43^. The steroids may also up-regulate myocylin gene (MYOC/GLC1A) expression, accumulation of which is an initial step to glaucoma^44, 45^. A number of other genes are thought to contribute as well though their significance are still being evaluated^46-48^. The elevated pressure in turn is transmitted posteriorly inducing a cascade of events that ultimately cause RGC axonal damage followed by RGC loss, mimicking the major etiology of primary open angle glaucoma (POAG).

While our study found an early elevation in IOP, it did not result in significant change to the RNFL thickness within the first 7 weeks in our study. The VN ratio, however, progressively decreased over this time period, suggesting that ultrastructural changes may precede NFL loss in early glaucoma. This finding/result corroborates with other preclinical studies having shown that RGC loss precedes RNFL thickness change following acute optic nerve injury ^49, 50^. As the VN ratio is derived from the superficial 30µm NFL around ONH, it primarily represents signal from RGC axons, in which microtubules contribute a significant portion of the measured reflectance^51^. Therefore, the decreased VN ratio may reflect the ultrastructural alterations in the microtubule ^52, 53^ leading to RGC damage caused by ocular hypertension. Indeed, previous studies by Huang *et al.* rigorously established that the loss of RGC cytoskeletons in axons <300 nm is an early event of RGC damage, and consequently leads to a “flatter” reflectance spectrum from RNFL^24^. This alteration which would yield a similar lower VN ratio as we found. This further consolidates our hypothesis that the decreased VN ratio would be a sensitive ultrastructural optical marker for early detection of RGC damage. Comparing to those previous studies that relies on the *ex vivo* measurement, vnOCT can be applied *in vivo* and isolate the signal from RNFL, allowing implementation in human eyes for clinical translation.

The alterations in the retinal circulation are considered important events in glaucoma development^54^. Lower ocular perfusion pressure and low systolic and diastolic blood pressure are known to be major risk factors for glaucoma^55^. Numerous studies have consistently shown the reduced blood flow^56-59^ in glaucomatous eyes. Recent studies using optical coherence tomography angiography (OCTA) have shown that the peripapillary capillary vasculature density (CVD) and capillary flow index are compromised in advanced glaucomatous eyes^10-15^, suggesting microvascular perfusion around ONH is an important factor in glaucoma pathogenesis. As retinal function relies on oxygen metabolism, changes in blood hemoglobin oxygen saturation (sO_2_) represent an attractive potential indicator of functional impairment. In our study, blood flow shows only a moderate decrease in dexamethasone group comparing to control group, at week 4 and 7. The increases of arteriovenous sO_2_ difference (A-V sO_2_), with decreases of the venous sO_2_, is more significant, indicating more oxygen extraction from the retinal circulation. Even with the mild decrease in blood flow, overall oxygen consumption from the retinal circulation increases, suggesting a state of oxygen hypermetabolism. This finding is in contrast to later stages of glaucoma when A-V sO_2_ is found to be decreased, due to advanced cell loss and decreased oxygen consumption ^60^. Yet, the retinal oxygen hypermetabolism is consistent with previous studies on an early stage diabetic retinopathy mouse model, which hypothesized increased oxygen extraction as a result of inflammation ^61^. The underlying biology is not clear at this moment and requires further investigation. Nevertheless, our data herein suggest that in early stage glaucoma, global alterations of sO_2_ might be another sensitive marker for early glaucoma.

The limitation of this study is the low number of data towards the end of the study, with four animals and five eyes per group producing good quality images and vnOCT readings, which affects the statistical power in Fig. 5C and 5D. Several animals were lost during the study, due to fighting injury; and some eyes had poor corneal quality, preventing reliable retinal imaging. Luckily, when we compiled the longitudinal data, the statistical power was sufficient to support the significant changes of VN ratio and AV-sO_2_ difference in dexamethasone group as compared to controls in ANOVA analysis.

## 5. CONCLUSION

In summary, we demonstrate that vnOCT can show ultrastructural changes in addition to changes in the oxygen levels in the retinal circulation in a dexamethasone-induced ocular hypertension mouse model. The changes were identifiable and seen even within the short-time frame of this study, indicating that pressure elevations can result in intracellular changes relatively quickly as well as possibly precede RNFL changes. These findings pave the way for future studies in mice and humans to gain further insight as to how changes progress, if they are reversible, and how they may respond to treatment.

## Supporting information

## ACKNOWLEDGMENT

The work contained in this paper has been supported by Bright Focus Foundation (G2017077), BU-CTSI 1KL2TR001411. We would like to thank Prof. Haiyan Gong, and Dr. Ruiyi Ren for their generous help on the dexamethasone-induced ocular hypertension mouse model and intraocular pressure measurement.

