## Supplementary material for "Early detection of retinal alteration by visible and near-infrared optical coherence tomography (vnOCT) in a dexamethasone-induced ocular hypertension mouse model"

### SUPPLEMENTAL FIGURES

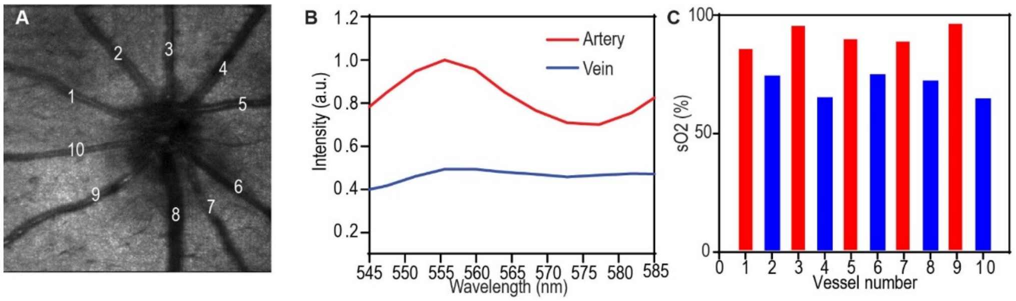


*Supplemental Fig. 1. An example of $sO_2$ calculation. (A) The en face projection from visible light channel. All major vessels were labeled. (B) Two example spectra extracted from blood vessel bottom wall for vessel 9 and 10. (C) The calculated $sO_2$ for each individual vessel. The arteries and veins are usually arranged alternatively, and easy to separate. The $sO_2$ reading for each retina averaged all the arteries and veins, respectively.*

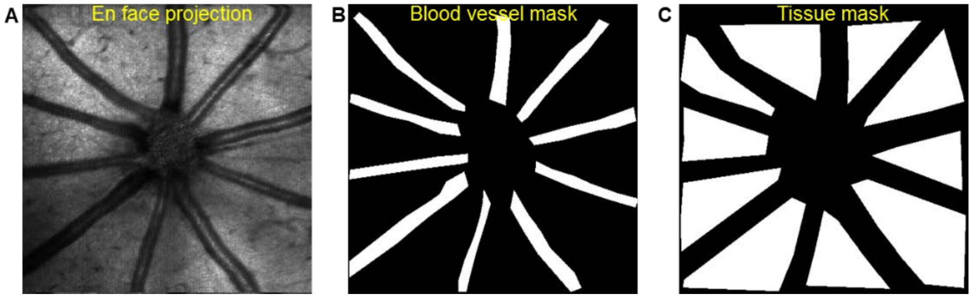


*Supplemental Fig. 2. An example of the ROI selection for blood vessels and retinal tissue. (A) The en face projection from visible light channel. (B) The segmentation of blood vessels for $sO_2$, and blood VN ratio calculation. (C) The ROIs for RNFL VN ratio calculations. Signals from all ROIs were averaged in visible light and NIR channels to calculate VN ratio.*
